# Acute kidney injury-induced circulating TNFR1/2 elevations correlate with persistent kidney injury and progression to fibrosis

**DOI:** 10.1101/2023.04.03.534558

**Authors:** Akshayakeerthi Arthanarisami, Yohei Komaru, Charikleia Katsouridi, Julian Schumacher, Deborah K. Verges, Liang Ning, Mai M. Abdelmageed, Andreas Herrlich, Eirini Kefaloyianni

## Abstract

**Background:** Elevated levels of circulating Tumor-Necrosis-Factor-Receptors 1 and 2 (cTNFR1/2) predict CKD progression. Whether acute kidney injury drives cTNFR1/2 elevations and whether they predict disease outcomes after AKI remains unknown.

**Methods:** We used AKI patient serum and urine samples, mouse models of kidney injury (ischemic, obstructive, toxic) and progression to fibrosis, nephrectomy, and related single cell RNA-sequencing datasets.

**Results:** We show that TNFR1/2 serum and urine levels are highly elevated in all mouse models of kidney injury tested, beginning within one-hour post-injury, and correlate with its severity. Consistent with this, serum and urine TNFR1/2 levels are increased in AKI patients and correlate with severity of kidney failure. Interestingly, the extracellular vesicle (EV)-bound forms of cTNFR1/2 correlate with renal function better than their soluble forms. TNF neutralization does not affect early cTNFR1/2 elevations, suggesting that cTNFR1/2 levels do not reflect injury-induced TNF activity. Kidney tissue expression of TNFR1/2 after AKI is only mildly increased and bilateral nephrectomies lead to strong cTNFR1/2 elevations, suggesting release of these receptors by extrarenal sources. cTNFR1/2 remain elevated for weeks after severe kidney injury and at these later timepoints cTNFR1/2 correlate to remaining kidney injury. During AKI-to-CKD transition, kidney expression of TNFR1/2 and cTNFR2 levels, correlate with development of fibrosis.

**Conclusions:** Our data demonstrate that AKI drives acute increases in cTNFR1/2 serum levels which negatively correlate with kidney function, in particular their EV-bound forms. Sustained TNFR1/2 elevations after kidney injury during AKI-to-CKD transition correlate with persistent tissue injury and progression to kidney fibrosis.

**Significance Statement:** Circulating TNF-Receptor-1 and -2 (cTNFR1/2) elevations predict chronic kidney disease progression. It remains unknown whether kidney injury can drive cTNFR1/2 elevations, whether cTNFR1/2 levels predict disease outcomes after AKI, and which are their circulating molecular forms. This study shows that kidney injury strongly increases cTNFR1/2 levels and that their extracellular vesicle-bound forms better correlate with kidney function loss, compared to their soluble forms. Sustained cTNFR1/2 elevations correlate to remaining kidney injury, and cTNFR1/2 levels post injury predict progression to kidney fibrosis. This work, thus, points to novel mechanisms for the observed cTNFR1/2 elevations in kidney disease and identifies unique predictive and diagnostic value in cTNFR1/2 elevations in AKI or AKI-to-CKD transition.

## Introduction

Since initial reports that circulating Tumor-Necrosis-Factor-Receptors 1 and 2 (cTNFR1/2) elevations predict diabetic nephropathy progression^1,2^, numerous studies have demonstrated that cTNFR1/2 level increases correlate with severity and progression of CKD of any cause examined (e.g., in additional diabetic nephropathy populations, idiopathic membranous nephropathy, IgA nephropathy, lupus nephritis and hypertension related nephropathy^3–12^). Elevated cTNFR1/2 levels also correlate with mortality in End-Stage-Renal-Disease patients and with nephropathy, cardiovascular events and mortality in diabetes^13–15^. Despite their clinical importance as markers of kidney disease progression, very little is known about the mechanisms of their release.

The transmembrane forms of cellular TNFR1 and TNFR2 can be cleaved by the transmembrane metalloprotease A-disintegrin-and-metalloprotease 17 (ADAM17), and their extracellular domains can be released as soluble proteins^16–19^; these constitute one part of cTNFR1/2. In addition, some of the cTNFR1 detected represents full-length protein on the surface of exosome-like vesicles^20^. It is not known whether this is also true for cTNFR2 and in which forms either of these receptors circulate in human kidney disease.

AKI episodes can lead to CKD development and/or enhance CKD progression. However, it remains unclear if acute kidney injury alone can cause cTNFR1/2 elevations. cTNFR1/2 levels have been measured to date either in CKD patients or related to AKI episodes in critically ill patients^21,22^ and in patients with pre-existing CKD^23^. Thus, whether cTNFR1/2 levels are elevated by acute kidney injury or during AKI-to-CKD transition remains unknown.

In this study, we examined the effect of AKI on cTNFR1/2 levels in different mouse models of disease, as well as in AKI patients without CKD or sepsis, and performed extracellular vesicle (EV) preparations to determine the contributions of soluble versus EV-bound forms of cTNFR1/2 to the total cTNFR1/2 pool. We also examined potential cTNFR1/2 cellular sources and the relationship of cTNFR1/2 with sustained kidney injury after AKI and with progression to fibrosis (AKI-to-CKD).

## Methods

### Human samples

Deidentified human samples of patients with AKI without CKD, sepsis or multiorgan failure, of patients with CKD or of healthy controls were provided by the Kidney Translational Research Core at Washington University in St. Louis. Cause of AKI was as follows: 15 acute tubular necrosis (ATN), one oxalate crystal ATN, one AKI of obstructive etiology, one contrast nephropathy, one ureterovesical junction stone AKI, one AKI with HIV, one AKI with malignant hypertension, one acute interstitial nephritis, one AKI with atypical hemolytic uremic syndrome, one AKI with hepatitis C-associated glomerulonephritis. AKI patients did not have clinical evidence of other organ failure based on clinical data review and serum biochemistries. CKD patient diagnosis were: four diabetic nephropathy (stage 3), three IgA nephropathy (stage 3), three FSGS (one stage 3, two stage 4) and three obstructive nephropathy (stage 3).

### Animal Experiments

8–12-week-old male mice were provided by Jackson Laboratories and were used in accordance with the animal care and use protocol approved by the Institutional Animal Care and Use Committee of Washington University School of Medicine, in adherence to standards set in the Guide for the Care and Use of Laboratory Animals. C57Bl6/NJ mice (used in all experiments in this study, unless noted) and FVB/N mice (used to confirm results in one experiment). Kidney ischemia for 18-27 min, as described in the text, at 37°C was induced bilaterally using the flank approach ^24^. A group of mice was subjected to unilateral ischemia (left kidney) for 25 minutes. Sham operations exposed kidneys without induction of ischemia. UUO was executed as described previously^25,26^. Briefly, after flank incision, the left ureter was tied off at the level of the lower pole with two 3.0 silk ties. Sham-operated mice underwent the same surgical procedure except for the ureter ligation.

Acute aristolochic acid nephropathy was induced in C57Bl6/NJ mice by a one-time IP injection of aristolochic acid (5, 10 or 20 mg per kg body weight, as described in the text, Sigma-Aldrich) in PBS. The control mice were administered the same amount of PBS. Cisplatin nephropathy was induced in C57Bl6/NJ mice by a one-time IP injection of cisplatin (20 mg per kg body weight, Sigma-Aldrich) in PBS. The control mice were administered the same amount of PBS. Murine etanercept (Pfizer) 12.5 mg/ml in PBS was injected into FVB/N mice at 10 mg per kg IP 2 h before surgery^27^. The control mice were administered the same amount of PBS. Anti-TNF antibody (MP6-XT22, eBioscience-Invitrogen) was diluted in PBS and injected into animals at 10 mg per kg IP 2 h before surgery. The control mice were administered the same amount of control IgG in PBS.

### Renal function

Serum creatinine was measured by LC-mass spectrometry at the O’Brien Core Center for AKI Research (University of Alabama School of Medicine). BUN levels were measured by DIUR-100 kit (BioAssay Systems).

### ELISA

Circulating TNFR1 and TNFR2 were measured in human or mouse samples using enzyme-linked immunosorbent assay (ELISA) kits (DY255, DY726, DY425 and DY426 respectively, all from R&D Systems) as per the manufacturer’s instructions. Serum dilutions for ELISAs were 100-fold for mouse and 50-fold for human samples. Urinary measurements were corrected for urinary creatinine, measured by the DICT-500 kit (BioAssay Systems).

### Extracellular vesicle preparations

Human and mouse serum samples were sequentially centrifuged for 20 min at 1,200 × g, and then for 35 min at 10,000 × g, followed by 90 min at 100,000 × g (pellets represent the exosome fraction; TLA100 rotor, Beckman-Coulter) and for 16 h at 175,000 × g (pellets represent the smaller particle fraction; TLA100 rotor Beckman-Coulter)^20^. The pellets from the last two centrifugations (separately or combined) and the supernatant from the last centrifugation (represents the soluble fraction) were used in ELISA, as described above.

### Single cell RNA sequencing

#### (i) mouse

scRNAseq analysis of four pooled sham or four pooled AKI kidney samples, was performed as described previously REF^28^. Briefly, cells were stained with propidium iodide, and live cells were sorted using FACSAria III (BD Biosciences). Libraries were prepared using the Chromium Single Cell 5’ Library Kit v2 and Chromium instrument (10x Genomics, Pleasanton, CA). Full-length cDNA was amplified, and libraries were submitted to Genome Technology Access Center of Washington University in St. Louis for sequencing at a depth of 50,000 reads. All processing steps were performed in R (v4.1.0) using Seurat v3^29^. Quality control was first performed on each library to find appropriate filtering thresholds for each. Expression matrices for each sample were loaded into R as Seurat objects, retaining only cells that have more than 200 and less than 3200 genes. Poor quality cells with >10% mitochondrial genes were removed. Any gene not expressed in at least three cells was removed. SCTransform was used for normalization, scaling, and variance stabilization (https://github.com/ChristophH/sctransform). This was done to reduce bias introduced by technical variation, sequencing depth, and capture efficiency. Integration of kidney single-cell data was done using Harmony to control for batch effects ^30^. After quality control and integration, 13,882 kidney cells were further analyzed. We identified 15 clusters by applying a K-nearest neighbor graph and clustering with a resolution of 0.2 on the result of principal component analysis (PCA). We visualized the clustering using uniform manifold approximation and projection (UMAP). To assign cluster identities, we first compiled a list of kidney cell types and their currently established markers ^31,32^ and performed manual annotation by those markers and additional differentially expressed genes (DEGs) between clusters identified by FindAllMarkers() function in Seurat. The expression levels of *Tnfrsf1a* (TNFR1) and *Tnfrsf1b* (TNFR2) in each cluster were visualized using ‘plot1cell’ package ^33^.

#### (ii) human

scRNAseq dataset from human kidney biopsy samples was obtained from the Kidney Precision Medicine Project (KPMP) website in Seurat object format in March, 2022 (https://www.kpmp.org/). After excluding the chronic kidney disease (CKD) cohort, this publicly available dataset included 20 participants in healthy reference group and 12 in AKI group. We performed cell-type annotation following their original labeling in the project, while excluding degenerative cells with marked loss of differentiation markers and/or increased %endoplasmic reticulum (ER) or %mitochondrial genes. Clusters with less than 10 cells in either of healthy or AKI group were removed. The expression levels of TNFRSF1A (TNFR1) and TNFRSF1B (TNFR2) by cluster were visualized in the aforementioned way.

### qPCR

Total RNA was isolated from mouse kidneys (whole kidney or cortex) using the Trizol (Invitrogen) following the manufacturers’ instructions. Total RNA was reverse transcribed using the QuantiTect RT Kit (QIAGEN) and real-time polymerase chain reaction (PCR) was performed with Fast SYBR Green (QIAGEN). Gapdh was used as the housekeeping gene. Primer sequences are provided in Suppl. Table 1. Data were analyzed using the ΔΔCt method.

### Statistics

All results are reported as the mean ± SEM. Comparison of 2 groups was performed using an unpaired, 2-tailed t-test or a Pearson correlation analysis where appropriate. Comparison of 3 or more groups was performed via ANOVA and Tukey’s post hoc test. Statistical analyses were performed using GraphPad Prism 9.0 (GraphPad Software Inc.). A *P* value of less than 0.05 was considered significant.

## Results

### Circulating TNFR1/2 levels are increased in mouse models of AKI and correlate with kidney injury levels

To test whether kidney injury can drive cTNFR1/2 elevations, we first measured cTNFR1/2 levels in the ischemia-reperfusion-injury (IRI) mouse model. We found that severe bilateral IRI, confirmed by BUN elevations above 100 mg/dL at 24 h post injury (**Figure 1A**), results in significant elevations of cTNFR1 and cTNFR2 levels 24 h post-IRI, approximately by 10- and 5-fold over their respective baseline levels (**Figure 1B-C**). Increasing ischemic injury, induced by increasing ischemia time and confirmed by increasing BUN elevations, results in increasing cTNFR1/2 concentrations, up to a plateau level (**Figure 1D**). At earlier time points post-injury, cTNFR1/2 levels show gradual increase starting already at 1 h post-IRI (**Figure 1E-F**). To test if cTNFR1/2 elevations represent a universal response to AKI, we used the nephrotoxic models of cisplatin and aristolochic acid. We found that both treatments result in increased cTNFR1/2 levels that mirror the course of kidney injury as assessed by BUN elevations (**Figure 1G-H**). A moderate (10 mg/kg) and high (20 mg/kg) dose of aristolochic acid result in moderate or severe injury respectively (assessed by BUN) and of corresponding cTNFR1/2 elevations (**Figure 1H**), confirming that increased severity of kidney injury results in increased cTNFR1/2 elevations.

**Figure 1:**
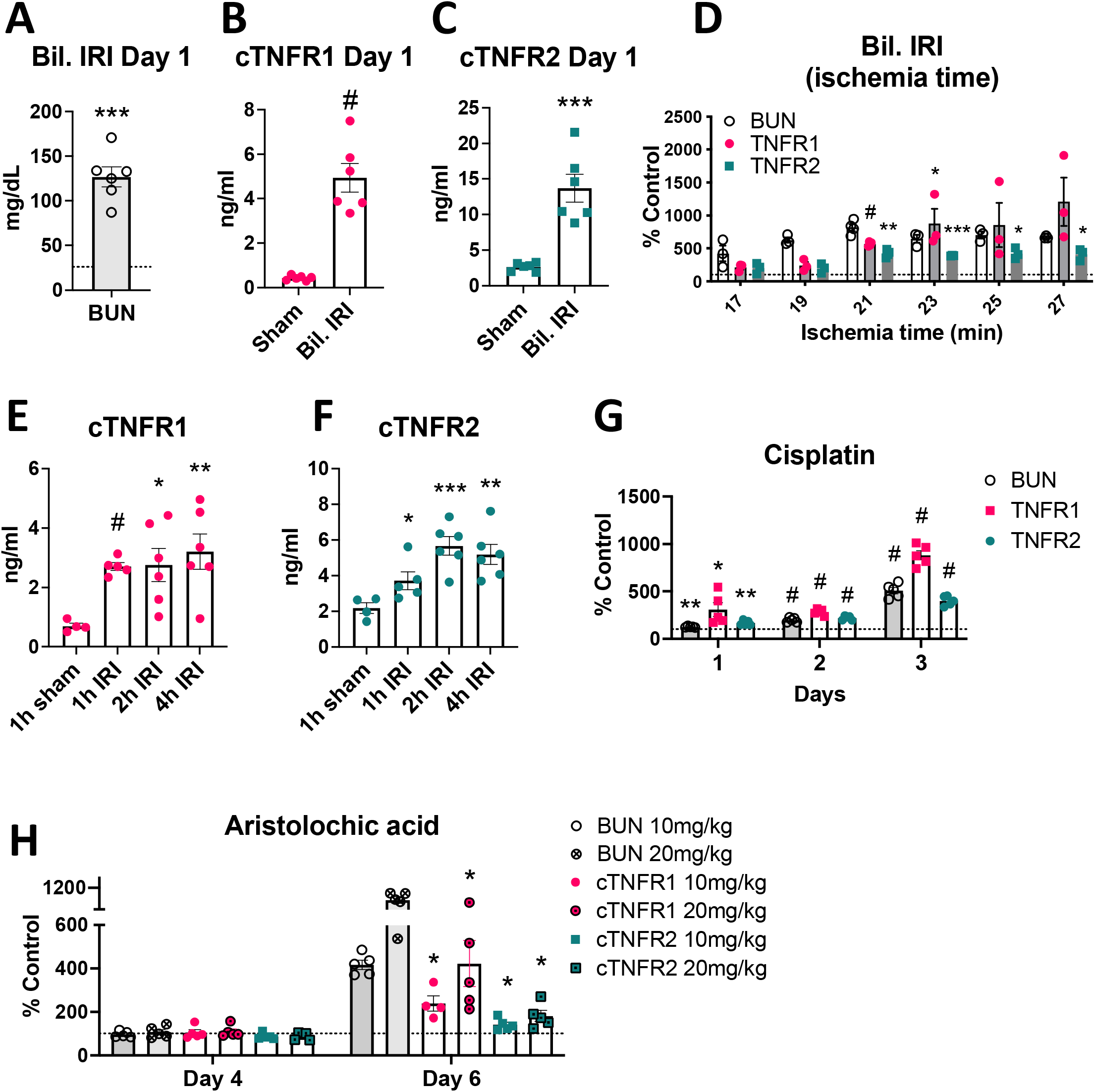
Circulating TNFR1/2 are increased in mouse AKI models and their levels correlate with kidney injury levels. **(A)** BUN levels at day 1 post injury in the severe bilateral IRI model (the dotted line denotes baseline levels). **(B)** cTNFR1 levels at day 1 post bilateral IRI or sham surgery. **(C)** TNFR2 levels at day 1 post bilateral IRI or sham surgery. **(D)** BUN, cTNFR1 and cTNFR2 levels at day 1 post bilateral IRI of increasing ischemia duration, expressed as percent over their respective baseline (% control) levels. **(E)** cTNFR1 levels measured at different time points (1-4h) post severe bilateral IRI or sham. **(F)** cTNFR2 levels measured at different time points (1-4h) post severe bilateral IRI or sham. **(G)** BUN, cTNFR1 and cTNFR2 levels measured at different time points (1-3 days) after cisplatin injection. **(H)** BUN, cTNFR1 and cTNFR2 levels measured at different time points (4 or 6 days) after aristolochic acid (10 or 20 mg/kg) administration. *: P<0.05, **: P<0.01, ***: P<0.001, #: P<0.0001

To determine whether the observed cTNFR1/2 increases were a result of reduced receptor clearance or of increased production/release, we tested TNFR1/2 urine levels in our AKI models. We found significant urinary TNFR1/2 elevations 24 h post bilateral IRI, that correlate with cTNFR1/2 serum levels (**Figure 2A-B**). We also tested cTNFR1/2 serum levels in unilateral kidney injury models, in which function in the uninjured kidney compensates for loss of function in the injured kidney. Both unilateral IRI and unilateral ureteral obstruction (UUO) result in significant elevations of cTNFR1/2, although BUN levels remain at baseline in both models (**Figure 2C-D**). These results suggest that cTNFR1/2 levels increase secondary to increased production/release rather than decreased receptor clearance.

**Figure 2:**
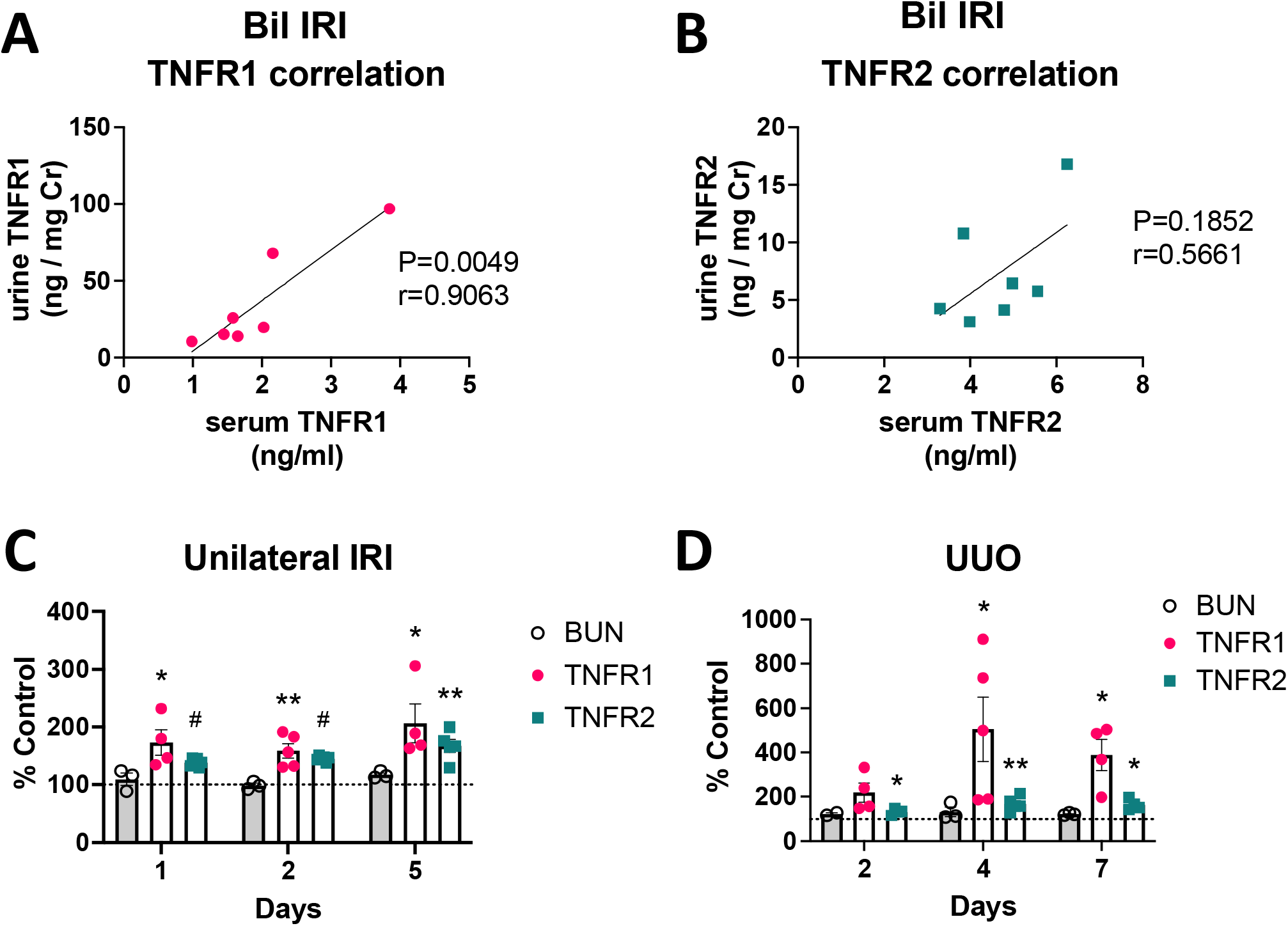
Urinary TNFR1/2 levels are increased by IRI, correlating to their serum levels, and cTNFR1/2 are increased in unilaterally kidney injury models. **(A)** Correlation of urinary TNFR1 levels with serum TNFR1 levels in matching samples, at day 1 post severe bilateral IRI. **(B)** Correlation of urinary TNFR2 levels with serum TNFR2 levels in matching samples, at day 1 post severe bilateral IRI. **(C)** BUN, cTNFR1 and cTNFR2 levels measured at different time points (1-5 days) after unilateral IRI. **(D)** BUN, cTNFR1 and cTNFR2 levels measured at different time points (2-7 days) after unilateral ureteral obstruction (UUO). Pearson correlation (r) value and P values are shown in (A) and (B). *: P<0.05, **: P<0.01

### Human AKI leads to increased cTNFR1/2 levels that correlate with AKI severity

We next measured the levels of TNFR1/2 in AKI patient samples. AKI induced significant upregulation of cTNFR1/2 serum levels, on average 4-5-fold over levels in healthy volunteers (**Figure 3A-B**). These serum elevations correlate with loss of kidney function, assessed by serum creatinine (**Figure 3C-D**). Similar to the mouse models, urinary TNFR1/2 levels are also elevated in AKI patients (**Figure 3E-F**) and correlate with corresponding serum levels (**Figure 3G-H**). These data show that human AKI results in significant cTNFR1/2 elevations that correlate with AKI severity, and in increased production/release of these receptors.

**Figure 3:**
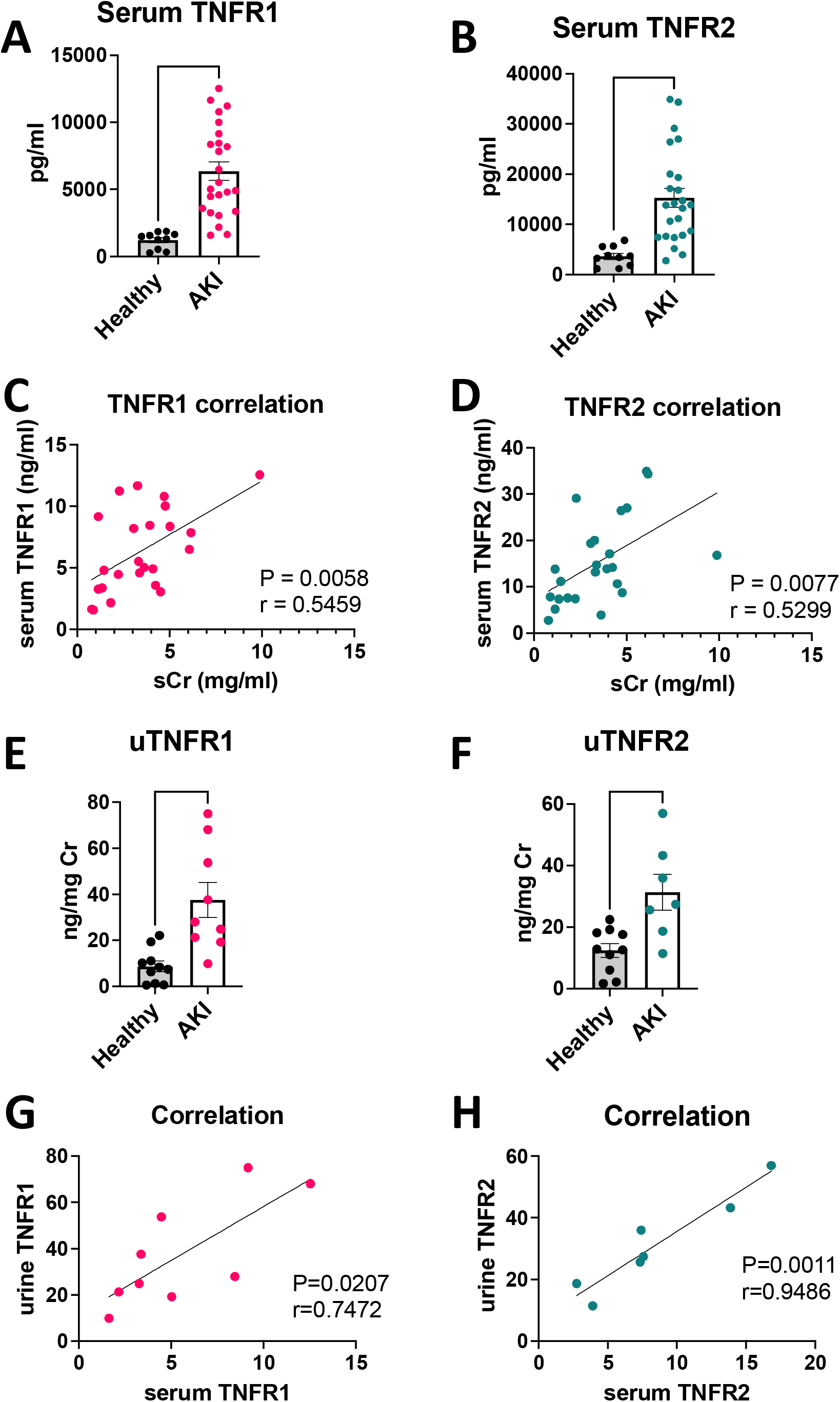
cTNFR1/2 are increased in human AKI and correlate with the severity of kidney failure. **(A)** Levels of cTNFR1 and **(B)** of cTNFR2 in healthy volunteer and AKI patient serum samples. **(C)** Correlation of cTNFR1 with serum creatinine in AKI patients. **(D)** Correlation of cTNFR2 with serum creatinine in AKI patients. **(E)** Urinary TNFR1 and **(F)** urinary TNFR2 in healthy volunteer and AKI patient samples. **(G)** Correlation of urinary TNFR1 to serum TNFR1 and **(H)** of urinary TNFR2 to serum TNFR2, in AKI patient samples. Pearson correlation (r) and respective P values are shown in graphs (C), (D), (G) and (H). **: P<0.01, ****: P<0.0001

### Extracellular vesicle-bound forms of cTNFR1/2 correlate with loss of renal function better than their soluble forms

cTNFR1 has been detected in exosome-like vesicles in human serum at baseline^20^ (we will refer to the combined exosome and smaller exosome-like vesicle-bound forms of the receptors as the “extracellular vesicle (EV)-bound forms”; **Suppl. Figure 1A**). Following the same method, we also detected cTNFR2 in the soluble and EV-bound fractions. Both the soluble and the EV-bound forms of TNFR1 and TNFR2 are increased in AKI patient serum by 4-5-fold on average, compared to healthy volunteers (**Figure 4A-B**). The ratio of soluble to EV-bound receptors does not significantly change between healthy and AKI patients (**Figure 4C-D**). Interestingly, however, the EV-bound forms of the receptors correlate much better with loss of renal function, assessed by serum creatinine measurements, compared to their soluble forms and, as a result, compared to their total levels (before EV-bound and soluble fractions are separated); this is demonstrated by improved Pearson correlations values (r values) and improved P-values (**Figure 4E-G**). EV-bound TNFR1 shows the highest correlation to serum creatinine in AKI. Similar results are obtained in a small number of CKD patients of various diagnosis (**Suppl. Figure 1B-D**). These results suggest that the EV-bound forms of cTNFR1/2 could provide a more accurate assessment of kidney function loss, compared to their total levels.

**Figure 4:**
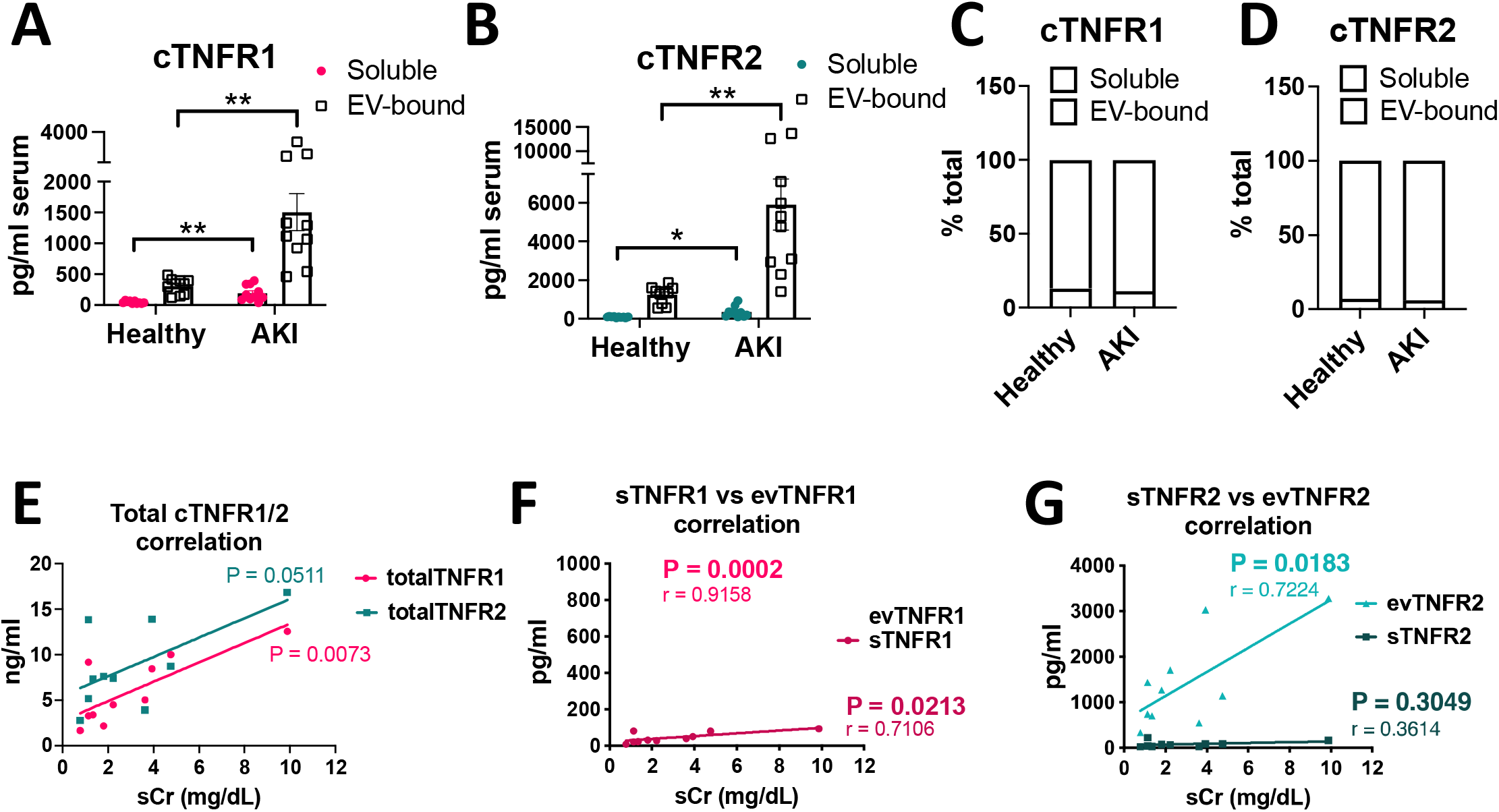
Extracellular vesicle-bound forms of cTNFR1/2 correlate to renal function better than their soluble forms in AKI patients. **(A)** Soluble and EV-bound forms of cTNFR1 from healthy volunteers or AKI patients, quantified by ELISA. **(B)** Soluble and EV-bound forms of cTNFR2 from healthy volunteers or AKI patients, quantified by ELISA. **(C)** Percentages of soluble and EV-bound cTNFR1 in healthy volunteers or AKI patients. **(D)** Percentages of soluble and EV-bound cTNFR2 in healthy volunteers or AKI patients. **(E)** Correlation of total cTNFR1 or cTNFR2 with serum creatinine. **(F)** Correlation of soluble and EV-bound cTNFR1 with serum creatinine. **(G)** Correlation of soluble and EV-bound cTNFR2 with serum creatinine. Pearson correlation (r) and respective P values are shown in graphs (E)-(G). *: P<0.05, **: P<0.01

### TNF neutralization does not affect early cTNFR1/2 serum elevations after kidney injury

It has been speculated, although not experimentally tested, that cTNFR1/2 are released as a result of cellular receptor activation by their ligand, TNF, representing markers of TNF activity in kidney or other diseases^34,35^. Indeed, TNF circulating levels are increased acutely after kidney injury, as seen in our AKI patients and in the severe bilateral IRI mouse model (**Figure 5A-5B**). To determine the contribution of TNF to cTNFR1/2 elevations in AKI, we neutralized TNF starting before injury and examined cTNFR1/2 at day 1 post IRI. We found that mice pre-treated with murine etanercept^27^ or an anti-TNF neutralizing antibody showed similar levels of kidney injury (**Figure 5C, 5E**) and similar cTNFR1/2 elevations (**Figure 5D, 5F**) compared to mice pre-treated with respective controls (cTNFR2 levels were not evaluated after etanercept (TNFR2-Fc fusion molecule) treatment because of its interference with endogenous TNFR2 detection). We thus conclude that signaling induced by TNF does not mediate early cTNFR1/2 elevations post AKI.

**Figure 5:**
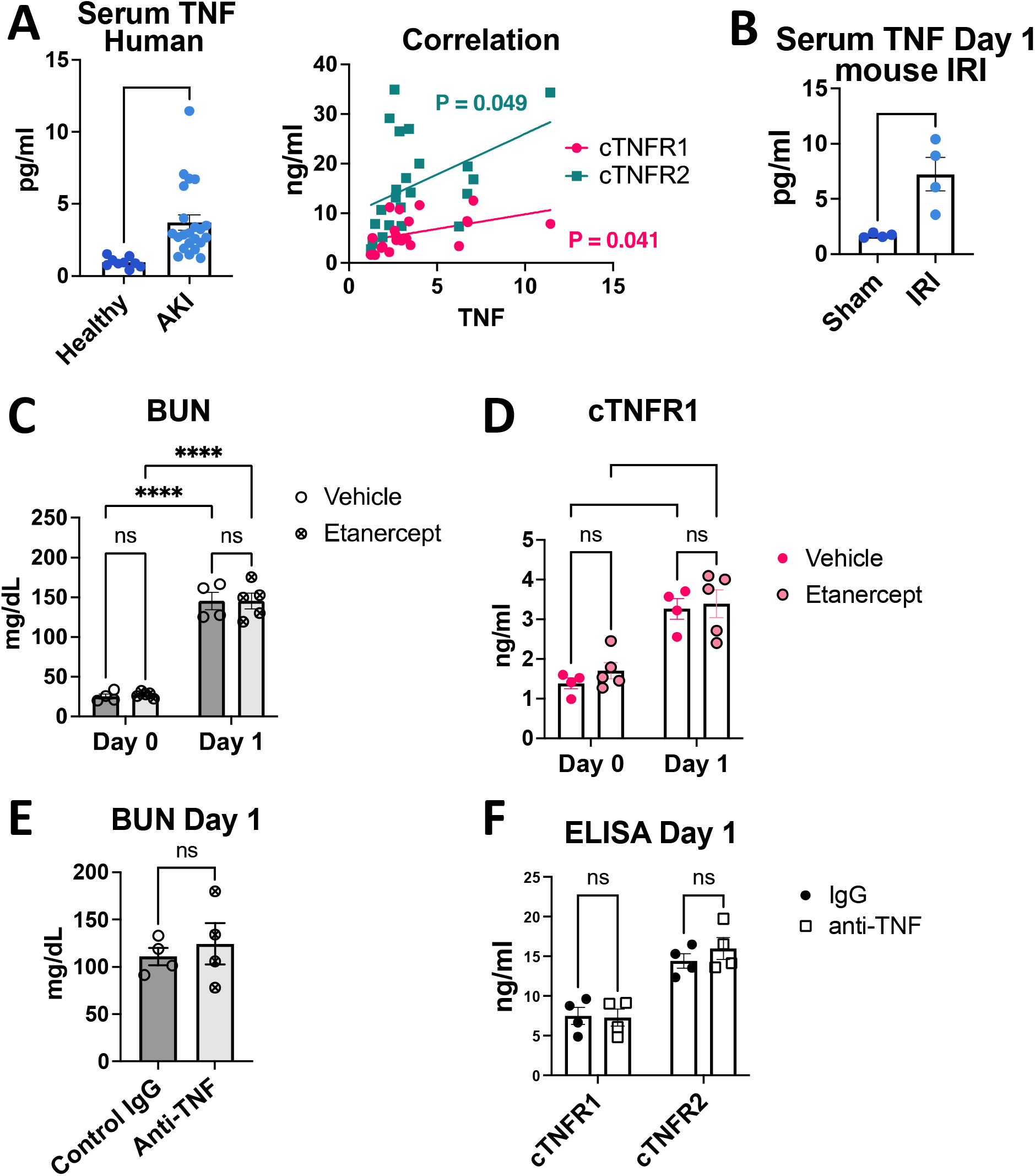
TNF neutralization does not affect cTNFR1/2 elevations early after bilateral IRI. **(A)** Serum TNF levels in healthy volunteers or AKI patients (left panel) and correlation to cTNFR1 and cTNFR2 in AKI (right panel). **(B)** Mouse serum TNF levels at day 1 post bilateral IRI or sham. **(C)** BUN levels at day 0 (before injury) and day 1 post bilateral IRI, in etanercept or vehicle-treated mice. **(D)** cTNFR1 levels at day 0 (before injury) and day 1 post bilateral IRI in etanercept or vehicle-treated mice. **(E)** BUN levels at day 1 post bilateral IRI in anti-TNF antibody or IgG-treated mice **(F)** cTNFR1 and cTNFR2 levels at day 1 post bilateral IRI in anti-TNF antibody or IgG-treated mice *: P<0.05, ****: P<0.0001, ns: not significant

### Early cTNFR1/2 elevations after AKI can be attributed to extrarenal sources

The injured kidney is a potential source of cTNFR1/2. To identify which kidney cell types express TNFR1 and TNFR2, we interrogated our existing single-cell RNAseq dataset of sham and IRI kidneys collected 24 h post injury^27,36^, a time point showing very strong elevations of serum and urine TNFR1/2 (**Figure 1B-C, 2A-B**). We found that TNFR1 (Tnfrsf1a) is ubiquitously expressed in the kidneys of sham mice, with the highest levels detected in endothelial cells (EC1 and EC2) and neutrophils (NT) (**Figure 6A**). Injury induced only mild upregulation of TNFR1 expression in ECs and very mild or no upregulation in other cell-types (**Figure 6A**). TNFR2 (Tnfrsf1b) is mostly expressed in immune cells, mainly in monocytes (Mono) (**Figure 6B**) and injury increases TNFR2 expression only mildly in monocytes and neutrophils (NT) (**Figure 6B**). We also examined TNFR1/2 kidney expression profiles in human AKI using single-cell analysis datasets from the Kidney-Precision-Medicine-Project (KPMP) atlas (https://atlas.kpmp.org/). The datasets include 20 healthy donor and 12 AKI patient samples. We found ubiquitous expression of TNFR1 (TNFRSF1A) in healthy donor samples and mild or no increases in AKI samples (**Figure 6C**). TNFR2 (TNFRSF1B) expression was detected mainly in immune cells, and overall decreased (with mild only increases in regulatory T cells (Treg) and some monocytes) in the AKI samples (**Figure 6D**).

**Figure 6:**
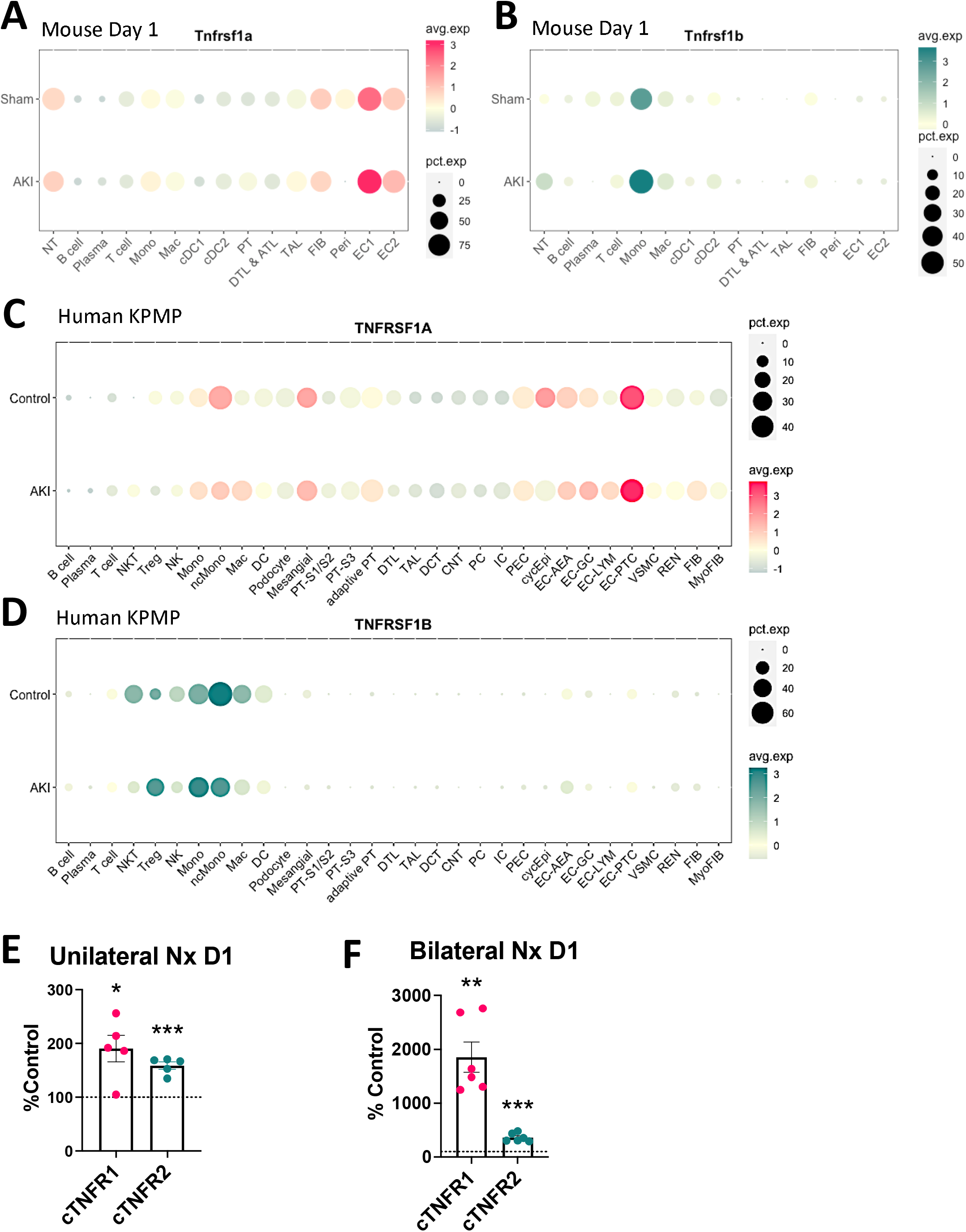
Kidney TNFR1/2 levels are mildly upregulated early after AKI while nephrectomy significantly increases circulating TNFR1/2. **(A)** Dot plot of TNFR1 (Tnfrsf1a) expression in mouse kidney cell types in sham and bilateral IRI at day 1 post injury. **(B)** Dot plot of TNFR2 (Tnfrsf1b) expression in mouse kidney cell types in sham and bilateral IRI at day 1 post injury. **(C)** Dot plot of TNFR1 (TNFRSF1A) expression in human kidney cell types from control and AKI patient samples. **(D)** Dot plot of TNFR2 (TNFRSF1B) expression in human kidney cell types from control and AKI patient samples. **(E)** cTNFR1 or cTNFR2 levels at day 1 post unilateral nephrectomy, expressed as percent of their respective baseline values (% control). **(F)** cTNFR1 or cTNFR2 levels at day 1 post bilateral nephrectomy, expressed as percent of their respective baseline (% control). Dotted line in (E) and (D) denotes baseline, *: P<0.05, **: P<0.01, ***: P<0.001

Considering the strong serum and urinary elevations we detect in mouse models and in human AKI (**Figures 1 and 3**), the above results suggest that cTNFR1/2 are released from existing receptors’ pools that do not require transcriptional upregulation and/or that they are released from extrarenal sources. To test cTNFR1/2 elevations after loss of kidney function and simultaneously eliminate kidney contributions to cTNFR1/2 levels, we utilized unilateral or bilateral nephrectomies. We found that unilateral or bilateral nephrectomy result in significant cTNFR1/2 increases after 24h (**Figure 6E-6F**). These results show that extrarenal sources can contribute significant amounts of cTNFR1/2 during loss of kidney function.

### cTNFR1/2 levels and their kidney expression levels remain elevated for weeks post kidney injury

Testing the long-term effects of kidney injury on cTNFR1/2 levels, we found that cTNFR1/2 remain elevated for at least four weeks post severe IRI (**Figure 7A**, C57Bl/6 mice; **Suppl. Figure 2A**, FVB/N mice), while traditional kidney function markers (such as BUN and serum creatinine) return to near baseline levels two to four weeks post-injury (depending on the initial injury and the mouse strain, **Suppl. Figure 2A**)^25,27^. We have previously reported elevations in kidney expression of TNFR1/2, beginning at day 2 post bilateral IRI, and at day 7 in the UUO model^25^. Analysis of a publicly available bulk RNA-seq^37^ dataset confirmed significant kidney TNFR2 expression increases at day 7 post bilateral-IRI that remained elevated for 12 months (**Suppl. Figure 2B**), while creatinine levels returned to normal at 14 days post injury in that study^37^. However, we found no significant correlation between cTNFR1/2 levels and kidney expression levels of each receptor (Tnfrsf1a and Tnfrsf1b genes respectively) at day 28 post severe IRI (**Figure 7B**), suggesting again that kidneys do not represent the major source of cTNFR1/2.

**Figure 7:**
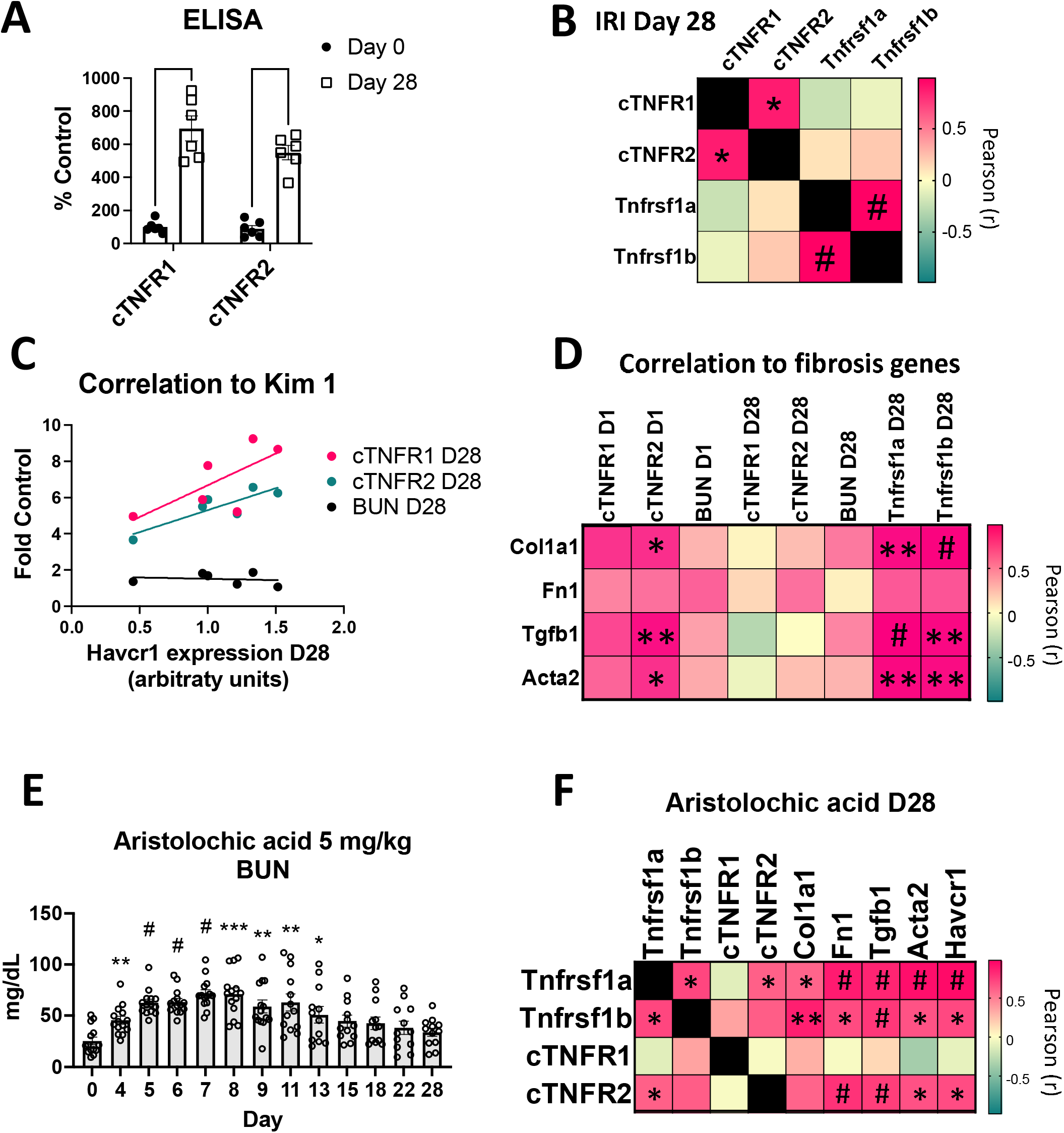
cTNFR1/2 and their kidney expression levels remain upregulated for weeks post injury and correlate with kidney injury and progression to fibrosis. **(A)** cTNFR1 and cTNFR2 levels, expressed as percent control (before injury), at 0 or 28 days post bilateral IRI (****: P<0.0001**) (B)** Pearson r value heatmap of correlations between cTNFR1, cTNFR2 and their respective kidney mRNA expression levels, at 28 days post bilateral IRI (*: P<0.05, #: P<0.001). **(C)** Correlations between cTNFR1, cTNFR2 or BUN (expressed as fold over their baseline levels, y-axis) with kidney Kim1 (Havcr1) mRNA expression levels (x-axis**). (D)** Pearson r value heatmap of BUN, cTNFR1 and cTNFR2 at day 1 and day 28 post IRI and of TNFR1 and TNFR2 (Tnfrsf1a and Tnfrsf1b genes respectively) kidney expression at day 28 post IRI with fibrosis marker kidney expression levels (*: P<0.05, **: P<0.01, #: P<0.001). **(E)** Time course of BUN levels in mice treated with low dose (5 mg/kg) aristolochic acid (*: P<0.05, **: P<0.01, ***: P<0.001, #: P<0.0001**). (F)** Pearson r value correlation heatmap of kidney expression levels of TNFR1 (Tnfrsf1a), TNFR2 (Tnfrsf1b), and of cTNFR1 and cTNFR2 with Kim1 and pro-fibrotic kidney gene expression, all at day 28 of the aristolochic acid model *: P<0.05, **: P<0.01, #: P<0.001).

### Sustained cTNFR1/2 elevations correlate with persistent injury and progression to fibrosis in AKI-to-CKD models

As a next step, we examined the hypothesis that cTNFR1/2 elevations correlate with AKI outcomes, in particular AKI-to-CKD progression and kidney fibrosis. Our severe-IRI model leads to development of fibrosis by three to four weeks post-injury, assessed by increased interstitial collagen accumulation and fibrotic-marker gene upregulation, and to loss of GFR which becomes detectable by 2-3 months post IRI^25,27,37,38^. Kidney injury molecule 1 (Kim1) is undetected in the healthy kidney and upregulated in injured proximal tubular cells after kidney injury and during transition to fibrosis. We found that cTNFR1/2 at day 28 post injury correlate with kidney Kim1 gene expression (Kim1 gene name: Havcr1) at the same time point, suggesting that sustained cTNFR1/2 levels indicate sustained tubular injury (**Figure 7C;** Pearson correlation r is 0.7030 and 0.8656 for cTNFR1 and cTNFR2 respectively, significant for cTNFR2: P=0.026). Of note, the traditional kidney injury marker BUN, shows no such correlation (**Figure 7C)**, suggesting that cTNFR1/2 levels can be very informative in identifying remaining kidney injury. Moreover, we examined the relationship of cTNFR1/2 levels with fibrosis markers expression levels (collagen 1α: Col1a1; fibronectin: Fn1; TGFβ: Tgfb1; α smooth muscle actin: Acta2) at day 28 post IRI (**Figure 7D**). cTNFR1/2 serum levels at day 28 do not correlate with those markers and neither does BUN levels (**Figure 7D, columns 4-6**). However, cTNFR1/2 serum levels at day 1 (in particular cTNFR2 levels) were predictive of subsequent fibrosis marker induction at day 28 (**Figure 7D, columns 1-2)**. Again, BUN at day 1 failed to show any correlation to kidney fibrosis at day 28 **(Figure 7D, column 3**). Of note, kidney TNFR1/2 expression levels (Tnfrsf1a and Tnfrsf1b respectively) are also increased at day 28 post injury and very strongly correlate with fibrosis markers (**Figure 7D, columns 7-8**), suggesting that the mechanisms that drive kidney TNFR1/2 expression and development of fibrosis are related.

In a different AKI-to-CKD model, low dose (5 mg/kg) aristolochic acid (AA) shows gradual elevations of BUN, which peak at Day 7 post AA injection (**Figure 7E**), and kidney fibrosis development at day 28 ^38,39^, a time point at which BUN has returned to baseline. Peak BUN (day 7) or day 28 BUN levels do not correlate with fibrosis gene expression at the end point, day 28 (**Suppl. Figure 2C-2D)**. Interestingly, we found significant correlation of cTNFR2 (but not of cTNFR1) serum levels at day 28 with the tubular injury marker Kim1 (Havcr1), with kidney fibrosis markers fibronectin (Fn1), TGFβ and α smooth muscle actin (Acta2), as well as a positive, although not significant, correlation with collagen 1α (Col1a1) expression (**Figure 7F**). Again, we see significant correlations of TNFR1 (Tnfrsf1a) and TNFR2 (Tnfrsf1b) kidney mRNA expression at day 28 with all fibrosis marker levels tested and with Kim1 expression (**Figure 7F**). Of note, there was again no correlation between kidney expression levels of TNFR1 and cTNFR1 and between kidney expression levels of TNFR2 and cTNFR2, suggesting that extrarenal tissues are the main circulating receptors’ sources at day 28 in the AA model.

Taken together, these results suggest that AKI-induced sustained TNFR1/2 serum elevations correlate with remaining kidney injury and that TNFR1/2 kidney expression and/or their circulating serum levels, particularly cTNFR2 levels, correlate with progression to kidney fibrosis.

## Discussion

In this study, we examined the effect of AKI on cTNFR1/2 levels. We found that kidney injury causes acute cTNFR1/2 increases, beginning within one-hour post-injury, with maximal levels around 24 h post-IRI. These elevations correlate to kidney function loss and are detected for prolonged periods, measured up to one month in our AKI-to-CKD mouse models. These results align with a recent report showing that CKD patients from the Chronic-Renal-Insufficiency-Cohort (CRIC) Study that had an AKI episode, showed increased cTNFR1/2 levels compared to CKD patients that did not have an AKI episode, and these elevations were detected months after the AKI episode^23^. It is intriguing that sustained cTNFR2, and to a lower degree cTNFR1, levels correlate with kidney Kim1 expression, an established and specific marker of tubular injury. Thus cTNFR1/2 elevations, in contrast to traditional markers of kidney function, such as BUN, are markers of persistent kidney injury during AKI-to-CKD transition. In addition, cTNFR2 levels (and to a lesser degree cTNFR1 levels) early after injury (day 1) correlate with kidney fibrosis markers levels during the transition to CKD (day 28). This is reminiscent of the predictive properties of cTNFR1/2 serum levels in CKD progression reported in numerous human studies. Based on the strong correlation of cTNFR1/2 levels to kidney function loss in AKI, their significant elevations in unilateral kidney injury models, their sustained upregulation in AKI-to-CKD progression, and their correlation with injury and fibrosis markers, we conclude that cTNFR1/2 serum levels represent sensitive markers of kidney function loss and underlying injury, outperforming BUN and serum creatinine. Whether this property is what renders them predictive markers of kidney disease progression in CKD or whether they also have disease modulating functions in AKI or CKD is not yet clear.

cTNFR2 is detected as soluble protein and in small exosome-like vesicles, as previously reported for cTNFR1^20^. We find that both soluble and EV-bound receptors are increased to a similar degree after AKI. This suggests some mechanistic connection between the release of soluble and EV-bound receptors. However, we find increased correlation of the EV-bound forms of cTNFR1/2 to the kidney functional biomarker serum creatinine, compared to their soluble forms (or their total levels), both in AKI and CKD serum samples. This finding suggests that there are some differences between EV-bound and soluble receptor release/clearance mechanisms that render the EV-bound forms better markers of kidney function. The result is very intriguing, considering the clinical importance of these molecules as kidney disease severity biomarkers. Longitudinal AKI or CKD patient studies are needed to examine if the EV-bound forms of the receptors also correlate better than their soluble forms to disease outcomes and progression.

TNFR1/2 kidney expression levels change only mildly 24 hours after injury in mice or in AKI patients, suggesting release by extrarenal sources. Although kidney TNFR1/2 expression is increased at later time points, receptors’ kidney expression does not correlate with their respective serum levels in any of our models, suggesting again extrarenal contributions. This is also corroborated by the fact that bilateral nephrectomies lead to very strong serum elevations of cTNFR1/2. These findings agree with a diabetic nephropathy study showing lack of correlation between circulating and kidney expression levels of TNFR1/2, using biopsies of patients obtained within six months before/after their cTNFR1/2 measurements^9^. Moreover, TNF neutralization does not affect early AKI cTNFR1/2 elevations, consistent with CKD studies that show correlation of cTNFR1/2 but not of TNF (total or free) with CKD progression^1,2^, suggesting different regulation mechanisms of the ligand’s versus the receptors’ serum levels. The exact cTNFR1/2 sources and release mechanisms require further investigation.

A potential explanation for the observed cTNFR1/2 serum elevations is the compromised circulating receptor clearance in kidney injury or after nephrectomy. This explanation is not likely the sole mechanism of cTNFR1/2 increases because, a) we detect significant elevations of TNFR1/2 in the urine in AKI, suggesting an increase of the total amount of receptors, and b) we detect increased cTNFR1/2 in models of unilateral injury, in which the remaining healthy kidney compensates for functional output, hence no increase in functional biomarkers (e.g., BUN, creatinine) is detected. This is also supported by CKD patients’ studies, which show increased levels of cTNFR1/2 and predictive value for disease progression even after correcting for eGFR, suggesting that their elevations are not merely a result of decreased clearance^1,2^.

Based on their significant predictive value in CKD progression and the fact that their elevations are detected long before development or progression of CKD, cTNFR1/2 could potentially have a causal role in disease progression. However, mechanistic studies exploring these functions in the context of in AKI, CKD or AKI-to-CKD transition have not yet been reported. An important impediment is the lack of knowledge of the molecular forms and release mechanisms of cTNFR1/2 in kidney disease. Our study provides novel insights into kidney injury-induced cTNFR1/2 elevations, that may enable future mechanistic studies, towards better understanding of cTNFR1/2 biology and potential roles in kidney disease progression.

## Acknowledgments

We would like to thank Pfizer for the provision of murine etanercept.

