## Supplementary material for "Acute kidney injury-induced circulating TNFR1/2 elevations correlate with persistent kidney injury and progression to fibrosis": Suppl Figures

**Supplemental Files**

Table of contents:

1. **Supplemental Figure 1:** Fractionation protocol and correlations of cTNFR1 and cTNFR2 forms with serum creatinine in CKD patient samples
2. **Supplemental Figure 2:** Long term TNFR1/2 elevations after injury and BUN correlations to fibrosis markers
3. **Supplemental Table 1.** qPCR primer sequences

Supplemental Figure 1: A. Fractionation protocol B-D. Correlations of circulating TNFR1 and TNFR2 with serum creatinine in CKD patient samples: B. Correlations of total levels of cTNFR1/2, C. Correlations of soluble and EV-bound forms of TNFR1, D. Correlations of soluble and EV-bound forms of TNFR2.

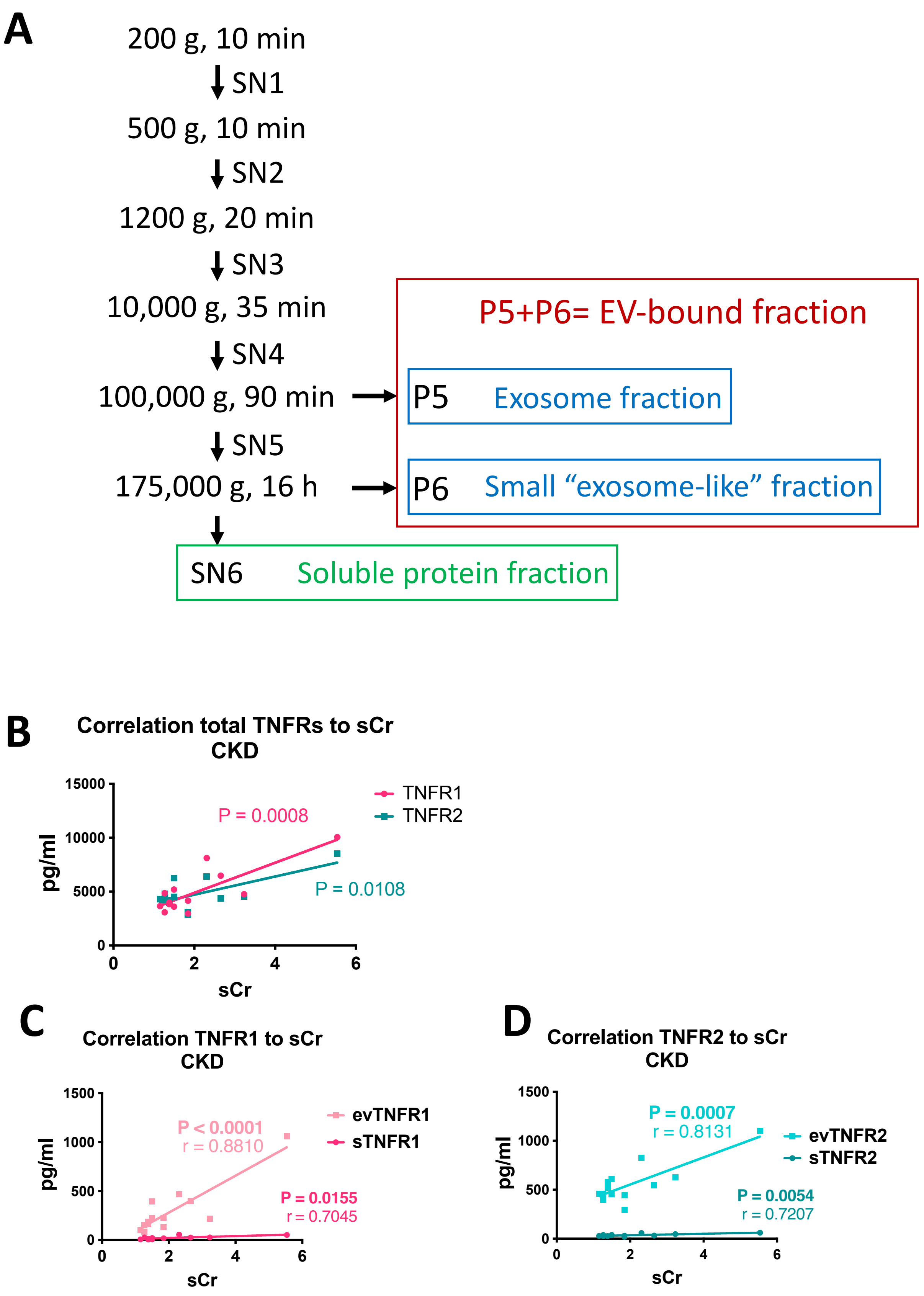

Supplemental Figure 2: Long term TNFR1/2 elevations after injury and BUN correlation to fibrosis markers. A. Sustained cTNFR1/2 elevations at day 28 post injury in the FVB/N mouse strain. B. Bulk mRNA sequencing analysis showing sustained kidney TNFR2 mRNA expression after IRI. C. Peak BUN (day 7) levels do not correlate to fibrosis/injury gene expression (day 28) in the aristolochic acid injury model D. BUN day 28 levels do not correlate to fibrosis/injury gene expression at day 28 of the aristolochic acid injury model.

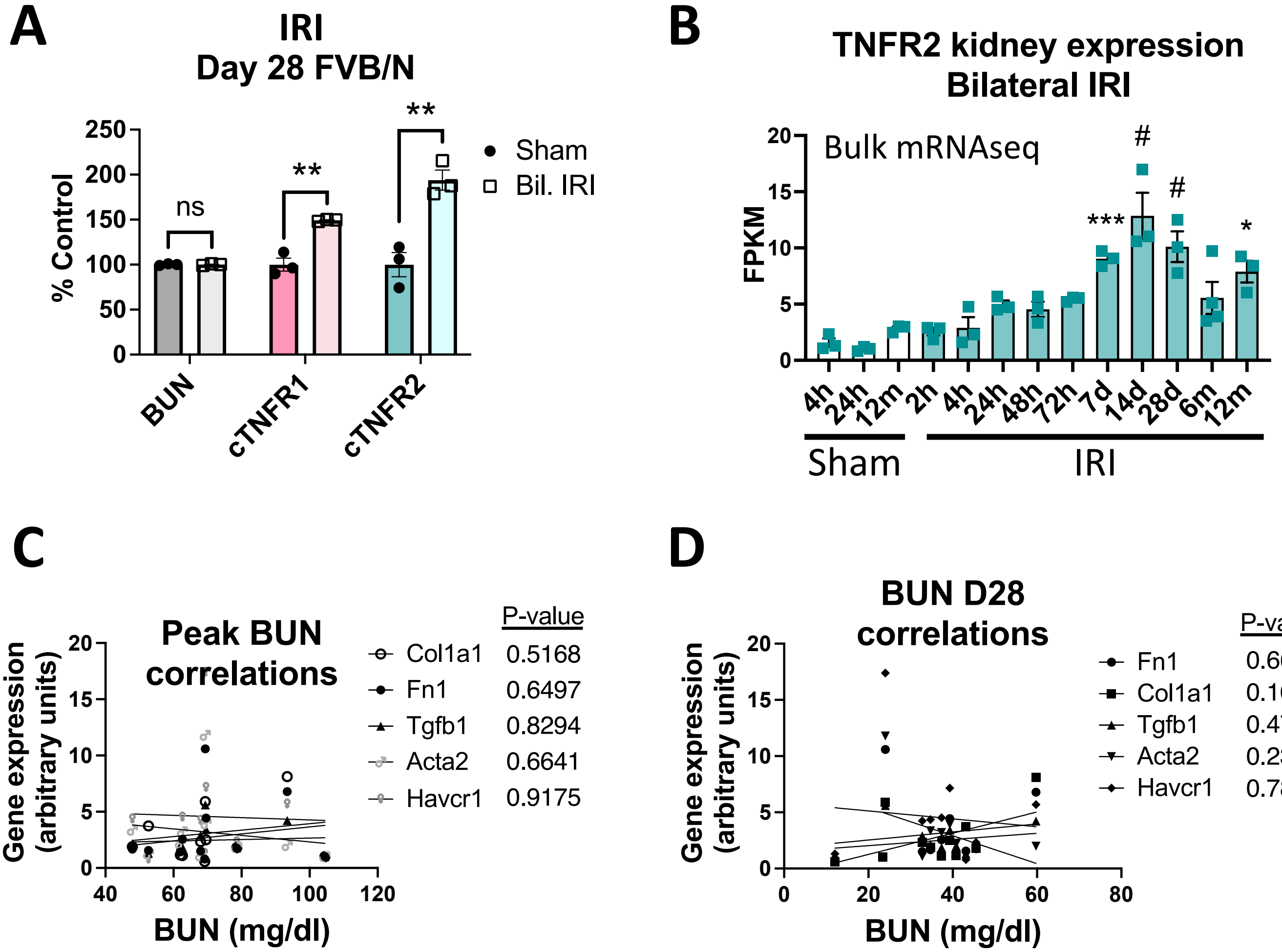

Supplemental Table 1. qPCR primer sequences

| Target gene | Synonym | Direction | Sequence |
|---|---|---|---|
| **Acta2** | αSMA | Forward | AGCCATCTTTCATTGGGATGGA |
| **Acta2** | αSMA | Reverse | TACCCCCTGACAGGACGTTG |
| **Col1a1** | collagen 1a | Forward | CCTGACGCATGGCCAAGAAG |
| **Col1a1** | collagen 1a | Reverse | TACCTCGGGTTTCCACGTCT |
| **Fn1** | fibronectin | Forward | GGTTCGGGAAGAGGTTGTGA |
| **Fn1** | fibronectin | Reverse | CACTCCTCTCCAATGGCGTA |
| **Gapdh** | | Forward | ACCACAGTCCATGCCATCAC |
| **Gapdh** | | Reverse | TCCACCACCCTGTTGCTGTA |
| **Havcr1** | Kim1 | Forward | ACAACAGCTGAGGTGACAGG |
| **Havcr1** | Kim1 | Reverse | TTCCCTGGAGGGATTGCTTC |
| **Tgfb1** | TGFβ | Forward | CTGCTGACCCCCACTGATAC |
| **Tgfb1** | TGFβ | Reverse | AGCCCTGTATTCCGTCTCCT |
| **Tnfrsf1a** | TNFR1 | Forward | ACGAATCACTCTGCTCCGTG |
| **Tnfrsf1a** | TNFR1 | Reverse | TCGCAAGGTCTGCATTGTCA |
| **Tnfrsf1b** | TNFR2 | Forward | GAGATGCCAAGGTGCCTCAT |
| **Tnfrsf1b** | TNFR2 | Reverse | AACTGGGTGCTGTGGTCAAC |
